# Memory reactivation in slow wave sleep enhances relational learning

**DOI:** 10.1101/2022.03.29.486197

**Authors:** Lorena Santamaria, Ibad Kashif, Niall McGinley, Penelope A. Lewis

## Abstract

Sleep has been shown to boost the integration of memories, and thus to facilitate relational learning. This benefit is thought to rely upon memory reactivation during non-REM sleep. We set out to test this possibility by explicitly cueing such reactivation using a technique called targeted memory reactivation (TMR), in which sounds are paired with learned material in wake and then softly played back to the participant in sleep, triggering reactivation of the associated memories. Specifically, we tested whether TMR during slow wave sleep leads to enhancements in inferential thinking in a transitive inference task. Because the Up-phase of the slow oscillation is more responsive to external cues than the Down-phase, we also asked whether stimulation at this specific phase is more beneficial for such integration. Our data show that Up-phase TMR boosts the ability to make inferences, but only for the most distant inferential leaps. Such stimulation was also associated with detectable memory reinstatement, whereas Down-phase stimulation produced no such trace and led to below-chance performance. These findings demonstrate that cueing memory reactivation at the right time points in sleep can provide a direct benefit to difficult relational learning problems.

**Significance Statement:** Memory reactivation in sleep is thought to be important for integrative thinking. We examined this by explicitly cueing reactivation of a transitive inference task during slow wave sleep using Targeted Memory Reactivation (TMR). Because TMR at different phases of the slow oscillation has different impacts, we cued different hierarchies at Up and Down phases. Up-phase TMR enhanced inferential performance and was associated with classifiable memory reactivation. Conversely, Down-phase TMR lead to a short-term impairment in inferential thinking and no detectable reactivation. These findings provide the first evidence that TMR can boost transitive inference and is thus important for integration and reasoning.

## Introduction

Relational memory is the ability to integrate multiple sources of knowledge, infer indirect associations between stimuli, and make decisions when presented with novel situations (Dymond and Llewellyn 2019; Lerner and Gluck 2019). One example of such integration is transitive inference (TI), or the deduction of the rankings of non-adjacent members of a linear hierarchy which has been presented via exposure to adjacent pairs. In simpler words, knowing A>B and B>C can allow deduction that A>C in an A>B>C hierarchy. Despite being studied for many decades in humans (Bryant and Trabasso 1971) and numerous other species (Camarena et al. 2018; Grosenick, Clement, and Fernald 2007; Lazareva et al. 2020) the mechanisms for TI remain elusive (Holyoak and Lu 2021; Morgan 2017). Furthermore, the dependence of TI on sleep has only been investigated relatively recently by a seminal study (Ellenbogen et al. 2007), and its replication (Werchan and Gómez 2013), which demonstrated that sleep is beneficial to this task, but only for the most distant inference pairs. While its role in TI remains unclear, sleep is thought to facilitate the abstraction of gist from recent experiences (Durrant, Cairney, and Lewis 2013; Lutz et al. 2017), and the integration of these with prior knowledge (Hennies et al. 2016; Tamminen et al. 2010). Prior learning is spontaneously reactivated during sleep, and this is important for such memory consolidation (Rasch and Born 2013). Importantly, memory reactivation can be directly cued by re-administering sensory stimuli that have been paired with learned information using a technique known as targeted memory reactivation (TMR), see (Hu et al. 2020) for a meta-analysis.

Although some prior work has linked cognitive performance enhancement with rapid eye movement sleep (REM), e.g. (Nolan 2010; Cai et al. 2009; Walker et al. 2002), most studies have focused on non-rapid eye movement sleep (NREM) and particularly on slow wave sleep (SWS), see (Born, Rasch, and Gais 2006; Schouten et al. 2017; Oudiette and Paller 2013; Rasch and Born 2013) for reviews. The main rhythms of SWS are slow oscillations (SO) and sleep-spindles. SOs are low-frequency oscillations at 0.5-4Hz that reflect alternation between hyper-polarized neuronal down phases and depolarized up-phases (Steriade, Nuñez, and Amzica 1993). SOs drive transient sleep-spindles at 9-16Hz (Steriade 2006) which are strongly linked to reactivation (Rasch and Born 2013). An elegant study (H.-V. V. Ngo and Staresina 2022) recently showed that cueing in the SO up-phase elicited more spindles and less forgetting than cueing in the down-phase, strongly suggesting that up-phase cueing is more likely to facilitate memory consolidation.

Here, we set out to investigate how reactivation of a TI hierarchy during SWS, and particularly during SO up and down phases, influences the ability to make inferences. To this end, we adapted the experimental set-up from (Ellenbogen et al. 2007) to include three hierarchies so we could apply closed-loop TMR (CL-TMR) in a within-subject design. Thus, one hierarchy was stimulated during the up-phase (Up condition), another during the down-phase (Down condition), and the remaining hierarchy was not stimulated (Control condition). Because recent work has shown that TMR effects can continue to unfold over time (Rakowska et al. 2021; Groch et al. 2017), our participants performed a third behavioral test two weeks after the manipulation. In keeping with other studies (Ellenbogen et al. 2007; Werchan and Gómez 2013), we expected cued reactivation in sleep to benefit only the inference pairs. We also predicted a benefit in the Up condition but not in the Down condition, and we expected the benefits to last, or even increase, over time.

Our data show that TMR in Up is associated with classifiable memory reinstatement and better inferential reasoning, and that this is strongest in the most distant inference pairs. On the other hand, stimulation in Down produced no evidence of reactivation and led to a temporary inhibition of inferential reasoning that recovered after two weeks.

## Materials and Methods

### Participants

Thirty adults (10 males, mean age 27 ±3.72) participated in the overnight experiment. All of them with no-self reported history of neurological, sleep or motor disorders. All participants completed a screen questionnaire before selection, provided written informed consent, and were reimbursed for their time. The experiment was approved by the School of Psychology Ethics Committee at Cardiff University. Participants agreed to abstain from caffeine and alcohol during the study and for 24-hours before. From the thirty participants that completed the task, 10 were eliminated either because of technical problems (n=3) or because they did not have enough stable SWS (n=7) to perform the stimulations (we required 12 rounds, these participants were mostly in light sleep and N2 stage). From the 20 participants left, 3 could not finish Session 3 due to the pandemic.

### Materials

The behavioral tasks were presented in a quiet room, participants were comfortable seated in front of the computer and stimuli were presented using Matlab^©^, Psychtoolbox (Brainard 1997) and Cogent 2000 (www.vislab.ucl.ac.uk). Three types of visual stimuli were presented to the participants: female faces (Lundqvist, Flykt, and Ohman 1998), outdoor scenes (taken from the internet) and unusual objects (Horst and Hout 2016), see figure 1B. Each stimulus was easily distinguishable from the others within and between categories. All items were presented in grey scale and matched for luminance. Each image was associated with an exclusive sound, semantically congruent with the image as closely as possible (e.g. bike image with a bike bell sound).

**Figure 1.**
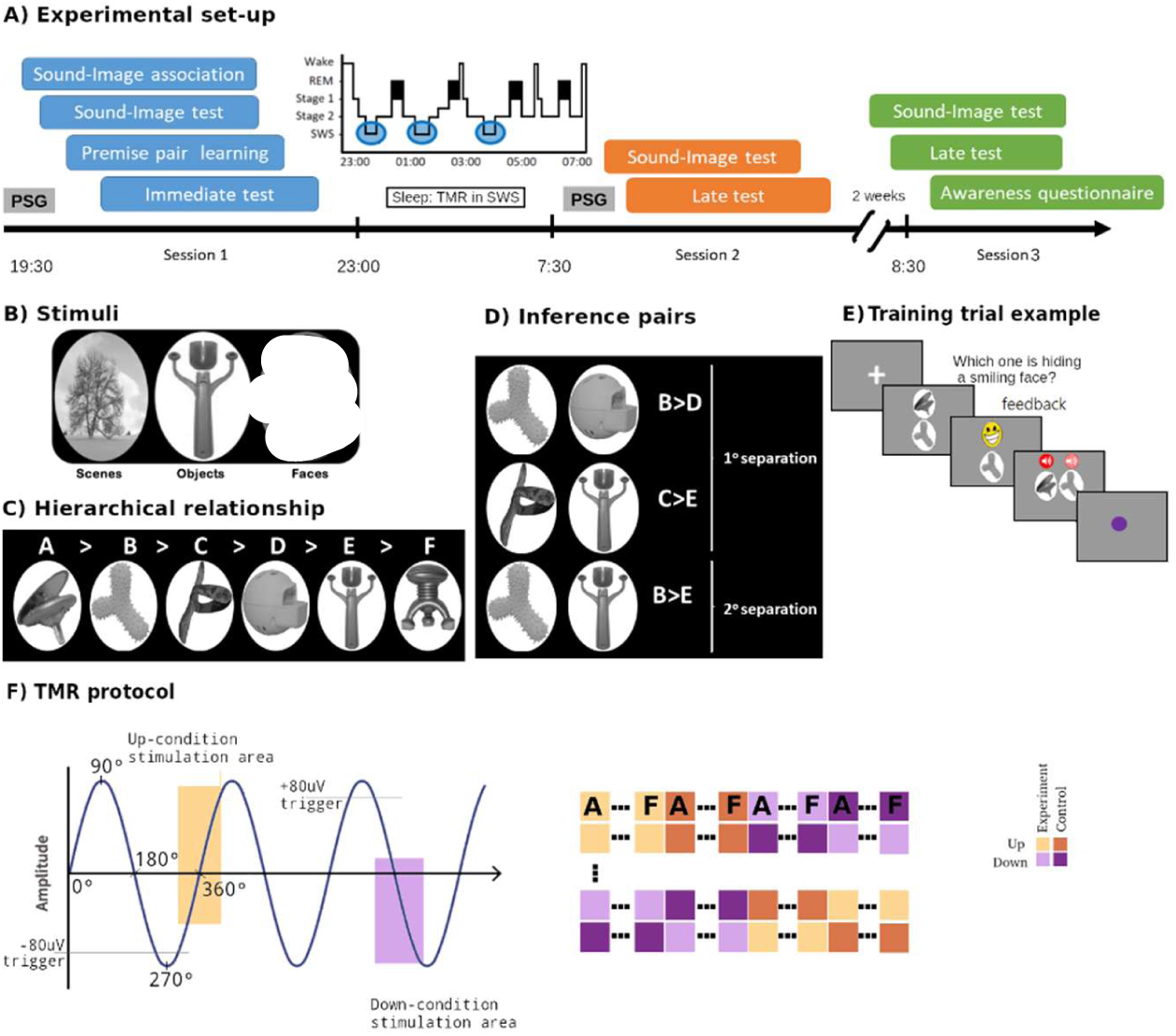
Experimental paradigm. A) Timeline: first, participants were wired-up (EEG). Participants in the first session completed four behavioral tasks: sound-image association, sound-image test, premise pair learning and immediate test. During learning, premise pairs of three hierarchies were presented. The Immediate test was the same as the learning counterpart but without feedback, allowing us to assess initial premise pair knowledge. Then, subjects went to sleep and TMR was carried out in SWS, where two out of the three hierarchies were stimulated. After waking up, still with the electrodes on, participants completed Session 2, which comprised a sound-image association test and a Late test. During the Late test, Inference pairs were presented for the first time, together with the previously learned premise pairs. After 2-weeks they returned to the lab for a third session, without EEG recording, where they completed the same tests as in Session 2 plus the awareness questionnaire. B) Example of the three categories used in the experiment: outdoor scenes, uncommon objects and female faces. C) Example of the visual stimuli arranged in a randomly determined hierarchical order. D) Resultant inference pairs from (C). E) Example of a typical learning trial. A cross appearing in the screen indicates the start of the trial (lasting between 0.5s to 1s), then two images of the hierarchy appear on the screen lasting until the user presses the up or down arrows. Feedback is provided immediately afterwards (a happy or angry face accordingly). To help sound-image consolidation, the sound of each stimulus is played while the stimulus is presented again on the screen. Finally, a purple dot is presented for 500ms to indicate the end of the trial. During the test part, the feedback and the sounds are eliminated. F) TMR protocol: left side the SO detection system used for the up (yellow) and down (purple) transition areas. On the right side and example of a “block”, where first were played the 6 items for the Up experimental condition, then Up-control sounds (new ones), followed by the Control and Experimental sounds of the Down condition respectively. Each element of the hierarchy was played in the right order for the experimental condition (A, B, ..F), additional control sounds were assigned to each hierarchy.

Sounds were taken from the internet and truncated into two different lengths, 2s and 200ms, and pitch normalized. We used the longer sounds in behavioral training, facilitating the sound-image encoding, and the shorter version for the rest of the behavioral tasks and TMR cueing. The sounds were played through noise-cancelling headphones (Sony MDR-ZX110NA) during behavioral tasks and through speakers (Dell A225) during sleep. The order of presentation of each stimulus category was counterbalanced across participants and the order of stimuli within each category was completely randomized for each subject. Hence, the experimenter was completely blind to which stimuli form the hierarchy and its order within each condition (Up, Down, Control), and which type of stimuli (faces, objects or scenes) was selected for each condition, so as not to influence the results. Each of the three hierarchies (figure 1B,C) comprised 6 images, each one with an associated (highly discriminable) sound. We prepared a set of 12 images and 12 sounds per hierarchy, that is a total of 36 images and sounds. At the beginning of the experiment, for each one of the three hierarchies, 6 of these images with their corresponding sounds were selected to be learned and the remaining 6 sounds were used as controls to be played during the TMR stimulation. Before participants started the first task, the 6 images which would be used during the experiment and the other 6 which would be used as control were randomly selected for each category (the experimenter was blind).

### Procedure

Participants arrived at the laboratory around 8 pm and changed into their sleepwear. They reported alertness by completing the KSS (Åkerstedt and Gillberg 1990) and SSS (Hoddes, Dement, and Zarcone 1972) questionnaires. Afterwards, they were fitted for PSG recording and performed the initial training and the immediate test explained in “Experimental tasks” section and figure 1A. Participants were ready for bed around 11pm. During the night, the previously learned tones were played softly during SWS. From the three stimulus categories, one was kept as a control (Control) and was not played during the night, allowing us to compare it against the other two which were cued during the Up and Down phases of the SOs respectively. After 7-8 hours of sleep, participants were woken at the agreed time and allowed > 20 min to overcome sleep inertia. During this time, they could go to the toilet, eat, and drink before completing the sleep quality, KSS and SSS questionnaires. Participants then completed the Late test and another Sound-Image association test (“Experimental tasks” section and figure 1A). Afterwards, the electrodes were removed, and participants could shower or go home. Finally, participants came back to the laboratory two weeks later (± 2days) to complete the second Late test and Sound-Image association test, identical to the previous one but without EEG recordings, to test the robustness of sleep-TMR mediated benefits.

### Experimental tasks by session

The experiment composed three sessions: evening (Session 1), next morning (Session 2) and a follow up session two-weeks after (Session 3). Each of the sessions was divided into different tasks as below:

a)Session 1: <u>Sound-image association learning task</u>: For each of the three categories, participants were shown each of the six items forming the category one by one. At the same time the associated sound (2s length) was played. Each sound-image pair was shown 4 times. The order of items within a category was randomized and the order of the categories themselves was counterbalanced across participants.

#### Sound-Image association test

Immediately after training, all participants performed a recall session to determine retention level. Three images were presented on the screen while a sound was played. Participants were asked to select as quickly and accurately as possible the image corresponding to the sound using the keyboard arrow keys. When they responded a rectangle surrounded the correct image (green if the participant’s selection was correct or red if it was wrong). Image screen position was randomized on every trial. The three images presented on the screen were pseudo-randomly selected, with the restriction that at least one of the two images was a ‘lure’ of the same category as the right answer. The sounds were cut down to only 200ms long. Participants performed two blocks with three repetitions of each sound per block. At the end of each block, accuracy was presented.

#### Premise pair learning task

Following previous related experiments (Ellenbogen et al. 2007; Werchan and Gómez 2013), all participants learned five relational premise pairs for each of the three categories. If each category formed a 6-item hierarchy, schematically represented as A>B>C>D>E>F (see figure 1C), the premise pairs would be: A>B, B>C, C>D, D>E, E>F. Where the notation “A>B” indicates “select A over B”. The pairs were presented one at a time, with images stacked vertically (figure1E). Subjects were instructed to select the item “hiding” a smiley emoticon from the two presented, at first by trial and error, but after practice and feedback they learned which item was correct. If they selected the correct item, it was replaced by a smiley emoticon. This is in line with (Werchan and Gómez 2013), where a smiling-emoticon was used as reinforcement. If they selected the wrong image it was replaced by an angry emoticon. After the feedback, participants received a second reinforcement as the pair was presented again but this time horizontally instead of vertically, and in the correct order (e.g. A-B) from left to right, with the corresponding sounds also played in the correct order. Pairs were organised into blocks of 10 trials for each of the hierarchies. This meant a total of 30 trials per block. Each block presented each of the five pairs of each hierarchy twice, counterbalancing the up-down positions (e.g. A above B and B above A, with A being the correct selection in both cases). The three hierarchies were not mixed within a block. For example, first all pairs for the “scenes” category were presented, then pairs in the “faces” category, and finally the “object” pairs. This order was counterbalanced across participants. Within each category, pairs were ordered pseudo-randomly to avoid explicitly revealing the hierarchy. Hence, a displayed pair cannot contain an item that was in the previous pair (e.g., A>B will never be followed by B>C). Furthermore, the order of the items within the hierarchy was randomly selected for each participant at the start of training, remaining unknown to the experimenter. At the end of each block, the overall performance for that block was shown on the screen to keep participants engaged with the task. All subjects underwent a minimum set of three blocks of training. After the third block, and every block thereafter, only performance of the “middle pairs”, meaning B-C, C-D and D-E, was saved to calculate the exit criteria (Werchan and Gómez 2013). If the averaged performance of these pairs for two of the last three blocks was over 66% for one of the hierarchies, the participant stopped receiving feedback for that hierarchy. However, all the premise pairs of this category still appeared on the screen to ensure the same number of trials/appearances for each hierarchy. This continued until the participant reached criteria for all the three hierarchies or a maximum of 10 blocks. In contrast to (Ellenbogen et al. 2007) and (Werchan and Gómez 2013), where the exit criterion was set to 75% accuracy for the middle premise pairs, we used a criterion of 66% to avoid ceiling effects and increase the chances of overnight improvement. On the other hand, we added a more restrictive criterion of 2 blocks out of 3 meeting the threshold, to be sure that the criterion was not achieved by chance. Similarly, to the above-mentioned studies, we only counted the middle premise pairs to evaluate the exit threshold because they are the necessary items for building the inferences.

#### Immediate test

After criterion was met, participants enjoyed a 5-minute break before proceeding to the

immediate test which assessed initial retention of the learned pairs. A similar protocol was used for testing and training with the exception that feedback and sound cues were removed in testing. Subjects were informed that they must select the right item based on previous learning. Participants performed four blocks, with 10 trials per hierarchy. Between blocks, participants solved arithmetic problems to clear short-term memory (von Hecker, Klauer, and Aßfalg 2019). Furthermore, pairs from the different hierarchies were randomly interleaved, always with the restriction of not showing the hierarchy explicitly.

b) Session 2: <u>Late test:</u> After filling the KSS and SSS questionnaires participants performed a similar test as before but this time, they were presented with previously learned premise pairs, new inference pairs, and one ‘anchor pair’ such that a total of 9 pairs were seen instead of just the 5 previously learned. The first new pairs were 3 inference pairs: B>D, B>E and C>E (see figure 1D). These pairs are named inference pairs because if you know that B>C and C>D, then you can infer that B>D. The inference pairs can be divided into 1st and 2nd degree of separation. That refers to the number of items between the ‘pair’ items, for instance between B and D is only one item (C), hence it has first degree separation, as does C-E. On the other hand, there are two items between B and E: C and D. Hence the B-E pair has a second degree of separation and is therefore the most distant pair within a 6-item hierarchy. Additionally, we also added a 4th pair, the anchor pair (A>F) as a control since inference is not needed to obtain this relationship. This is due to the fact that A is always correct and F is always incorrect (von Hecker, Klauer, and Aßfalg 2019). Participants were instructed that they might see novel combinations and if that was the case, they should try to make their best guess. At the end of each trial, they were confidence from -2 (guessing) to +2 (certain) using the up and down arrows. Following a similar protocol to Session 1, participants performed four blocks with math exercises between them.

#### Sound-Image association test

After a 5-minute break, subjects performed a new sound-image association test with the same structure as the Session 1’s test but without feedback.

c) Session 3: This used the same tasks in the same order as in Session 2. However, this time participants’ brain activity was not recorded. Finally, the participants performed an Awareness questionnaire (see figure 1-1).

### Closed-loop TMR protocol

The two categories that used for TMR and the control were counterbalanced across all participants (see SM3). The control category was not cued during the night. From the other two, one was assigned to Up and the other to Down. Stimulation started after participants entered stable SWS and was halted for arousals or any other sleep stage. Participants were exposed to an extra control-hierarchy of sounds for each of the two TMR conditions. These control-hierarchy-sounds, also 200ms duration, were completely novel to the participants and were included to allow us to distinguish the TMR effect from a normal brain ERP. Each one of the hierarchies, was composed of 6 items and played in order: A, B…, F. The order of both experimental and control hierarchies and of Up and Down cueing was randomized and counterbalanced across blocks. Each block comprised four hierarchies: Experimental Up, Control Up, experimental Down, control Down (see figure 1F). The minimum number of blocks to include a participant in the analysis was 12, this means 288 cues were presented during the night. Online detection of SOs was based on the detection of the negative half-wave peaks of oscillations. The electrode used as reference for the on-line detection was F3 as frontal regions are a predominant SOs area (Massimini et al. 2007), band-pass filtered in the slow-wave range (0.5-4Hz). When the amplitude of the signal passed a threshold of -80uV the auditory stimulus was delivered after a fixed delay of 500ms (H.-V. V. V. Ngo et al. 2013; H. V. V. H.-V. V. Ngo et al. 2015). Inter-trials intervals were set to a minimum of 4 seconds, that is after every sound played there was a minimum pause of 4s. The SO detection, auditory stimulation and presentation of the trigger to the EEG recording was via a custom-made Matlab-based toolbox (https://github.com/mnavarretem).

### EEG recordings

Sleep was recorded using standard polysomnography including EEG, electromyographic (EMG) and electrooculography (EOG). EEG was recorded using a 64-channel LiveAmp amplifier (Brain Vision^©^). Electrode impedance was kept below 10KΩ and sampling rate was 500Hz and referenced to Cpz electrode. In addition to the online identification of sleep stages, polysomnographic recordings were scored offline by 3 independent raters according to the ASSM criteria (Berry et al. 2015), all of them were blind to the periods when the sounds were reactivated.

### EEG analysis

Pre-processing and analysis were all performed with Fieldtrip (Oostenveld et al. 2011) and custom Matlab functions. Data were low-pass and high-pass filtered (30Hz and 0.5Hz respectively). Eye and muscle related artefacts were removed using independent component analysis (ICA). Bad channels were interpolated (spline interpolation) and data was re-referenced to linked mastoids. We calculated ERPs by segmenting the cleaned signal into 4 second segments, from -1s before stimulus onset to 3s afterwards. A final visual inspection of the dataset was performed, and any residual artefact was manually removed. To calculate the differences between stimulation conditions we averaged across all trials, but for the classification analysis we kept the trial information. To study the time frequency evoked TMR response, we calculated the power spectrum of the signal locked to the TMR cue onset using Morlet wavelets from 4 to 20Hz with 0.5Hz resolution and a time window from -1 to 2.4s in 50ms steps at the subject level. The width of the wavelet was set to at least 4 cycles per time window, adaptatively to the frequency of interest. Resulting TFRs were then expressed as the relative change of baseline from -1s to 0ms pre stimulus onset.

### Statistics

Statistical assessment of EEG data was based on nonparametric cluster permutation tests with the following parameters: 2,000 permutations, two-tailed, cluster threshold of p<0.05, and a final threshold of p<0.05 using Fieldtrip toolbox (Oostenveld et al. 2011). The time-frequency statistical analysis was restricted to the post-cue interval (0 to 2.4ms) to avoid the natural differences of the Up and Down phases of the SOs before cueing onset. To examine the accuracy of the TMR protocol (circular statistics) the R package Circular was used (Jammalamadaka, S. Rao and SenGupta 2001). Behavioral analysis was performed using robust statistical methods from the R package WRS2 (Mair and Wilcox 2020) to avoid any possible issues with normality and homoscedasticity assumptions. Repeated measures analysis of variance (RM-ANOVA) or simple 1-way ANOVA analysis was performed accordingly, always keeping the trial information and adding individual differences into the analysis (subjects ID’s). One sample t-test (Students or Wilcoxon signed-rank) tested for difference over chance level (50%) of each group, Condition and Session of interest. Significance of Pearson correlations between classification performance and behavior used a bootstrap method implemented in R, “boot” package (Buckland, Davison, and Hinkley 1998).

### Classification

Classification of single-trial data was performed using MVPA-light (Treder 2020) for each participant and each time point (−1 to 3s) using the sleep-ERP values (filtered between 4 to 20Hz) of the 60 EEG channels as feature. Performance of two classifiers was compared using a linear discriminant analysis (LDA) and a support vector machine with linear kernel (SVM). We used a 5-fold cross-validation method with 2 repetitions and principal component analysis (PCA) to reduce dimensionality (n=20). The data within each fold was z-scored to avoid bias. Additionally, we used two different metrics to evaluate performance of each classifier: traditional accuracy (ACC), defined as % correct predictions, and area under the curve (AUC), or trade-off between the true positive and false positive rates. Once classifiers were calculated for each participant, we performed a between-subject cluster analysis (Treder 2020) to determine at what time points the Experimental and Control sounds were statistically different for each condition. All code for the analysis of this study is available at https://github.com/Contrerana.

## Results

Prior to sleep, participants performed a transitive inference task (figure 1A) with three hierarchies (figure 1B) of 6-items each (figure 1C). Each of the items was associated to a separate semantically related sound (e.g., traffic noise with a city landscape). Participants learned the sound-image associations until they were over 90% accurate in a retrieval task. They then learned the transitive inference task by repeatedly viewing adjacent (premise) pairs (e.g. A-B), for each hierarchy, and indicating which hid the smiley face, with feedback (figure 1E). After a 5-minute delay, the same task was performed without feedback to assess learning level (immediate test). Next morning, participants performed this test again, but inference pairs were now included, B-D, C-E and B-E, (delayed test, figure 1D). Sounds associated with two out of the three hierarchies were presented during SWS, with one hierarchy presented in Up and another in Down using CL-TMR. To control for other EEG underlying mechanisms induced by the cueing, control sounds were also played during the night for both states, Up and Down.

## Behavioral results

### Premise pairs

#### Training Performance

Immediate test performance on premise pairs was above chance (50%) in all three conditions (Down: 74.7±0.02%, Control: 79.2±0.02%, Up: 79.6±0.02%) but not at ceiling (see table 2-2). A 1-way ANOVA showed no significant differences between these conditions at baseline (F=2.69, *p*=0.07). Further post-hoc analysis (2-tailed t-test) showed no differences: Down vs Control (*p*=0.095), Down vs Up (*p*=0.095), Up vs Control (*p*=0.93). Hence, participants had equal premise pair knowledge for all conditions in the immediate test. Performance on the sound-image association was over 90% for all the stimuli (figure 2-1) and the number of times each stimulus was used for each condition remained relatively constant, table 2-1. Hence, we can rule out any bias due to stimuli or cue-memory associations.

**Figure 2.**
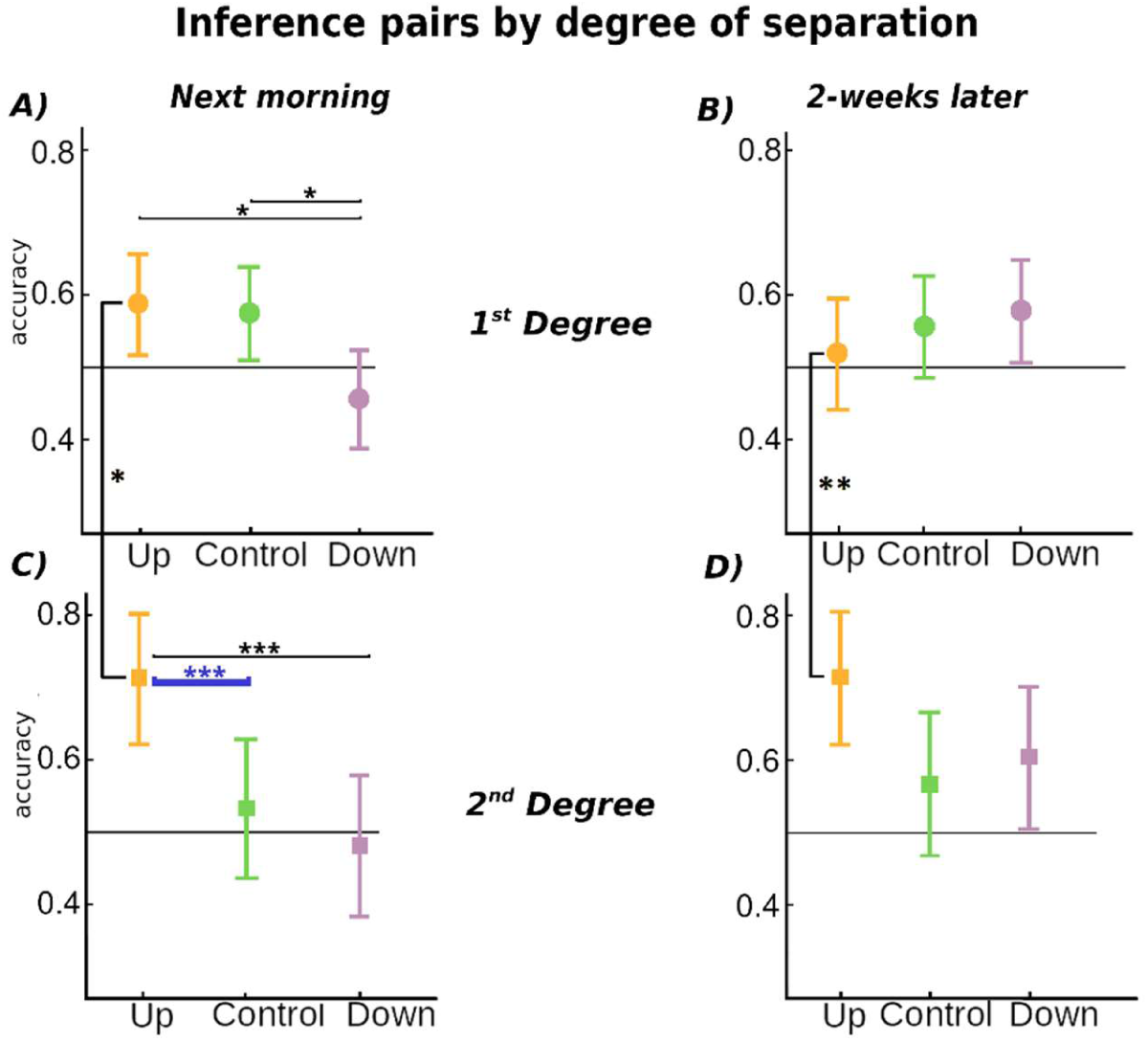
Behavioural results. Inference pair performance for each condition: Up (yellow colours), Down (purple) and Control (greenish). Inference pairs performance separated by degree of separation: A) 1^st^ degree of separation for Session 2 (next morning) and B) Session 3 (follow-up), C) 2^nd^ degree of separation for Session 2 and D) Session 3. Bars represents 95% confident intervals. Statistically significant differences are indicated as: * p<0.05, ** p<0.01 and ***p<0.001. All results are corrected for multiple comparisons. Blue line indicates statistically significant difference from Control condition.

To examine how in the premise pair performance evolved over time, and whether there were any impacts of TMR on this, we performed a repeated measures (RM)-ANOVA with Session (3 levels) and Condition (3 levels) as within subject factors, and with accuracy as the dependent variable (see Figure 2-2). We found main effects of Condition (F=3.5, p=0.029) and Session (F=32.39, p<0.0001), but no interaction (p=0.72). Further analysis for Condition revealed no significant differences between conditions in any of the three sessions (smallest *p*=0.103). On the other hand, there were clear Session effects: both between Session 1 (pre-sleep) and Session 3 (two-weeks later), and between Session 2 (next morning) and Session 3, figure 2-2 (all p<0.001). Thus, in keeping with normal declarative forgetting, there was a marked drop in premise pair performance over two weeks irrespective of condition, see table 2-2 for details. There was also no overnight improvement in premise pair accuracy, but this is in line with previous TI literature (Ellenbogen et al. 2007; Werchan and Gómez 2013). This result might be surprising given that associative memories are often strengthened by sleep (Rasch and Born 2013), but is also in line with the idea that sleep facilitates more weakly encoded memories (Schapiro et al. 2017), though this could also depend on other factors such as the type of task or the strength of the cue-memory associations (Denis et al. 2021).

### Inference pairs

Inference pairs were introduced in the Late test performed during Session 2 and repeated in the 2-week follow up (Session 3). To examine the effect of TMR on inference pair performance both the next day and two weeks later, and how this differed for close (1st degree) and distant (2nd degree) inferences, we performed a RM-ANOVA with the factors Session (2 levels), Condition (3 levels) and Degree of separation (2 levels) (see Figure 1D). This revealed main effects of Degree (F=5.91, *p*=0.016) and Condition (F=13.97, *p*=0.002) but not Session (F=1.91, *p*=0.17). This ANOVA also revealed two interactions: between Degree and Condition (F=9.89, *p*=0.008), and between Session and Condition (F=7.86, *p*=0.021), but no interaction between Session and Degree (*p*=0.601). Finally, the interaction between all three factors was not significant (F=0.65, *p*=0.722). We examine the significant results in further detail below.

#### Interaction between Degree and Condition

To directly investigate the interaction between Degree and Condition, our post-hoc t-tests collapsed across Session. This showed that the Up condition differed from both Control (*p*=0.015) and Down (*p*=0.017) at the 2^nd^ Degree only. There was also a difference between 1st and 2nd degree in the Up condition (*p*=0.004), but there were no differences between the Control and Down conditions for either Degree of separation (lowest *p*=0.79). All probabilities were corrected for multiple comparisons, see figure 2, 2-3 and table 2-3. These results clearly demonstrate that Up-phase TMR can provide a benefit to transitive inference, but in keeping with (Ellenbogen et al. 2007), this is only significant for the most distant items.

To examine how the effects of Condition on 2^nd^ degree items change over time we performed separate ANOVAs for 2^nd^ degree items at Sessions 2 and 3. At Session 2 this showed better performance for Up vs. Control (*p*=0.027) and Down (*p*=0.006), but not between Down and Control (*p*=0.880), all corrected for multiple comparisons, figure 2C and table 2-3. At Session 3, two-weeks later, comparisons of Up vs. Control and Up vs. Down were no longer significant (*p*=0.087 and *p*=0.251 respectively). The benefit of the Up stimulation therefore fades over the two-week consolidation period (Figure 2D).

In summary, stimulation of the SO up state was associated with significantly better performance on the most distant (2nd Degree) inference pairs compared to Control stimulation in the next-day data, figure 2B lower left panel. Interestingly, stimulation of the Down phase appeared to impair inferences when compared to Up, irrespective of degree of separation (figure 2, left panel). However, this picture changed after two-weeks, when performance on inference pairs stimulated in the Down phase improved to above chance levels and did not differ from performance on pairs in the Control and Up conditions (figure 2A, right panel). Performance on Up and Control conditions did not change over this retention interval.

#### Interaction between Session and Condition

Post-hoc analysis of the interaction between Session and Condition revealed a difference between Up and Down conditions (*p*<0.001, corrected for multiple comparisons) at Session 2, supporting the idea that the Up and Down states are associated with distinct forms of neural processing (H.-V. V. Ngo and Staresina 2022). However, comparison of Up and Down with Control showed no differences after correction for multiple comparisons (*p*=0.46 and *p*=0.19 respectively). No differences were found for Session 3. This analysis also revealed an unexpected difference between Sessions for the Down condition (*p*=0.038), with a significant improvement of performance from under chance level (50%) at Session 2 (46.4±2.8%) to over chance level two-weeks later (58.6±2.9%). Control and Up conditions remained over chance level across both sessions, figure 3A and tables 3-7 for full report.

**Figure 3.**
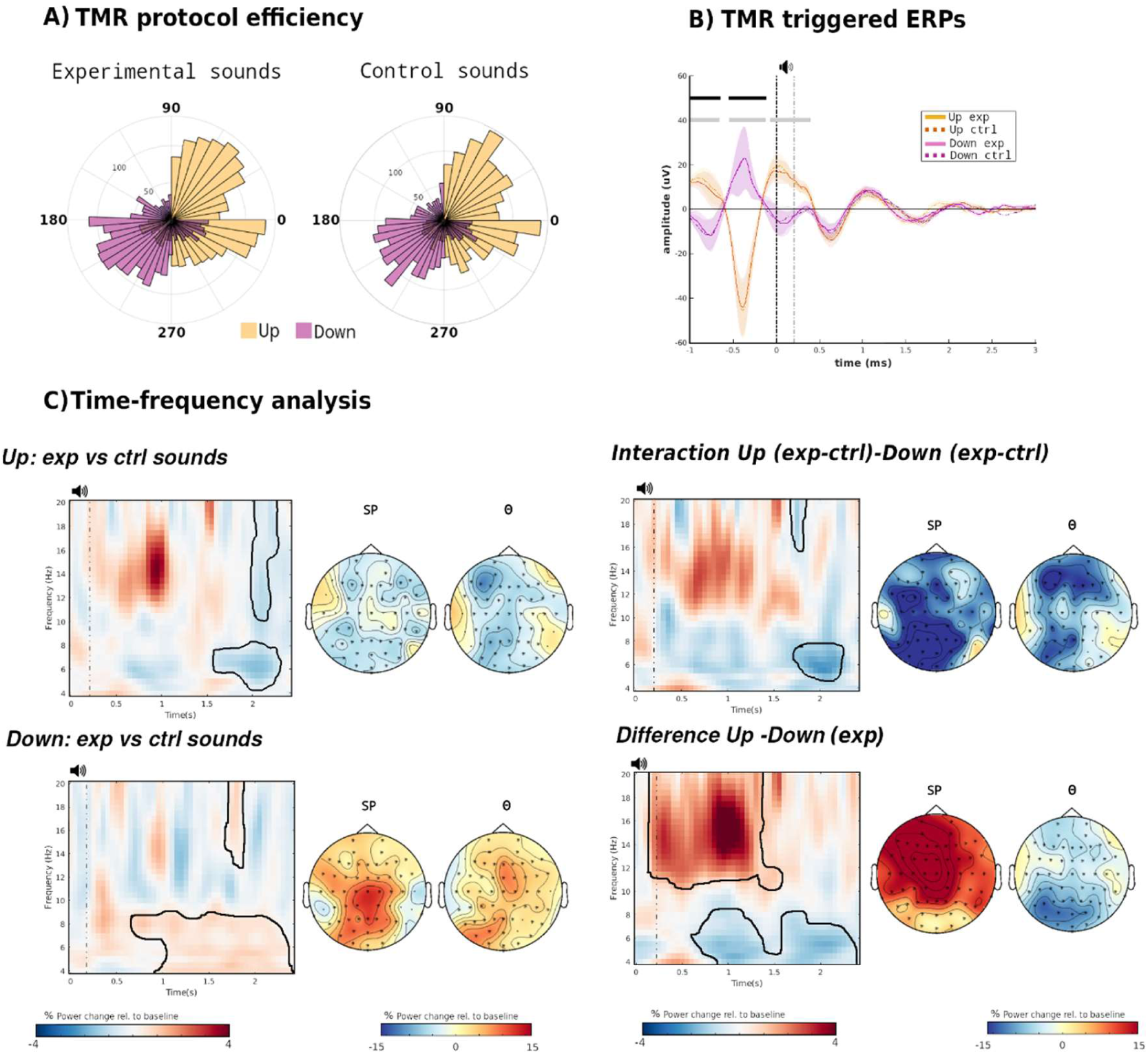
EEG analysis. A) Phase angle analysis at stimulus release per each trial. The left polar histogram represents the Experimental sounds with Up condition in yellow and Down condition in purple. Similarly, the right histogram represents the Control sounds. B) ERP statistics. ERPs for Up (yellow) and Down (purple) Conditions and Experimental (solid line) and Control (dashed line) sounds. All ERP-data are shown as mean±SEM (standard error of the mean) across participants. There are no differences between Experimental and Control sounds but there are differences, as expected, between Up and Down stimulations. These differences are highlighted in grey for the Control sounds and black for the Experimental sounds. All graphs are the values for electrode F3. C) Time-frequency analysis results (grand average at F3 channel). Experimental vs. Control (new) sounds power spectrum differences are plotted for Up condition (top left), Down (bottom left), their interaction (top right) and a direct comparison Up vs Down for Experimental sounds (bottom right). Vertical dashes lines indicate the onset of the auditory TMR cue (200ms). The black contour outlines significant clusters (two tailed, p<0.05). The power spectrum differences from those significant clusters were used to plot the topographies on the right side of each graph. Topographies were divided into theta (θ:5-8Hz, right side) and a general spindles frequency band (SP:10-20Hz, left side). Channels involved in the significant cluster are highlighted with an asterisk.

Global effect of TMR: Finally, because the 1st Degree items also showed a trend towards benefit from TMR, we collapsed across both Session and Degree for an exploratory analysis to test our *a-priori* hypothesis that Up state cueing would benefit all inferences. This showed better performance after Up than Control across all stimuli (*p*=0.03). Performance was also better for Up vs Down stimulation (*p*=0.002), but not for Control vs Down (*p*=0.687), all corrected for multiple comparisons.

### EEG results

To distinguish the effect of TMR upon EEG responses from the effect of a playing a sound, we used both Experimental sounds, which were associated with the previously learned hierarchy and Control sounds which were associated with an unlearned hierarchy as indicated in figure 1F.

#### Efficiency of the closed-loop algorithm

To check that our online algorithm correctly differentiated between the Up and Down conditions, we calculated event-related potentials (ERPs) separately for each Condition (Up/Down) and Cue type (Experimental/Control sounds) (figure 3B). In addition to the expected ERP differences between Up and Down before cue onset, there was a third significant cluster for Control but not Experimental sounds. Furthermore, after the sound offset, both cues and type of sounds elicited a second SO cycle that made them statistically indistinguishable in line with previous literature (Göldi et al. 2019; H.-V. V. Ngo and Staresina 2022). To corroborate the accuracy of our closed-loop algorithm we calculated the phase of the cortical SO at stimulus onset for each trial and participant (figure 3A and figure 3-1). For Experimental sounds, the average values at channel F3 were 358.20° (standard deviation (SD):0.58) and 205.61° (SD:0.44) for the Up and Down conditions respectively. Similar values were obtained for the Control sounds: 358.79° (SD:0.59) and 208.37° (SD:0.50) respectively. Circular statistics corroborate a significant difference between Up and Down conditions for both Experimental and Control (p<0.001) but no differences between Experimental and Control in either Down or Up condition (p>0.1).

Time-frequency analysis at the stimulation electrode (F3) compared Experimental vs. Control sounds for frequencies between 4 and 20Hz for Up and Down cueing. For Up, there was a significant power decrease in both, SP (cluster *p*=0.008, 2.05s to 2.1s) and theta bands (cluster *p*=0.008, 1.6s to 2.3s) towards the end of the trial, figure 3C. For Down, the opposite was observed, with a power decrease apparent (*p*=0.012, t=1.7s to 1.8s and *p*=0.010, t=0.7s to 2.35s). There also was a significant interaction between cueing Condition and type of sound for both SP band (p=0.077, t=1.7s to 1.8s) and theta (p=0.007, t=1.6s to 2.3s). These temporal late differences are in line with previous work (Göldi et al. 2019; Cairney et al. 2018). We also compared Up and Down directly for the Experimental sounds to assess the beneficial effect of the Up condition. There is a positive cluster (Experimental > Control) around the sound onset (p=0.008) for SP band lasting until around 1.25s. This is similar to other CL-TMR literature (H.-V. V. Ngo and Staresina 2022). On the other hand, for theta band there is a decrease in power (Experimental < Control) with a significant cluster (p=0.037) ranging from 0.65s to 2.4s.

#### Detection of reactivation

To determine if memory-related neural activity was reactivated during the night, we trained two machine learning algorithms to differentiate between Experimental and Control sounds for each condition (Up/Down). After calculating the classifiably of data for each participant, we performed cluster statistics at the group level for each condition (Treder 2020). Only the Up condition presented a significant cluster of above-chance classification. This cluster (p=0.037) ranges from 1204 to 1298ms after stimulus onset for the SVM classifier with area under the curve (AUC) as performance metric (figure 4A). Similar values were obtained for the rest of the tested combinations, tables 4-1 to 44 & figure 4-1. There were no significant clusters for the Down condition (lowest p=0.074). The absence of significant classification here could indicate that the brain is not able to activate the corresponding memory trace above chance when the cues are presented in the down phase. Hence, no behavioral benefit is obtained the following morning. However, this does not explain the performance improvement observed two weeks later.

**Figure 4:**
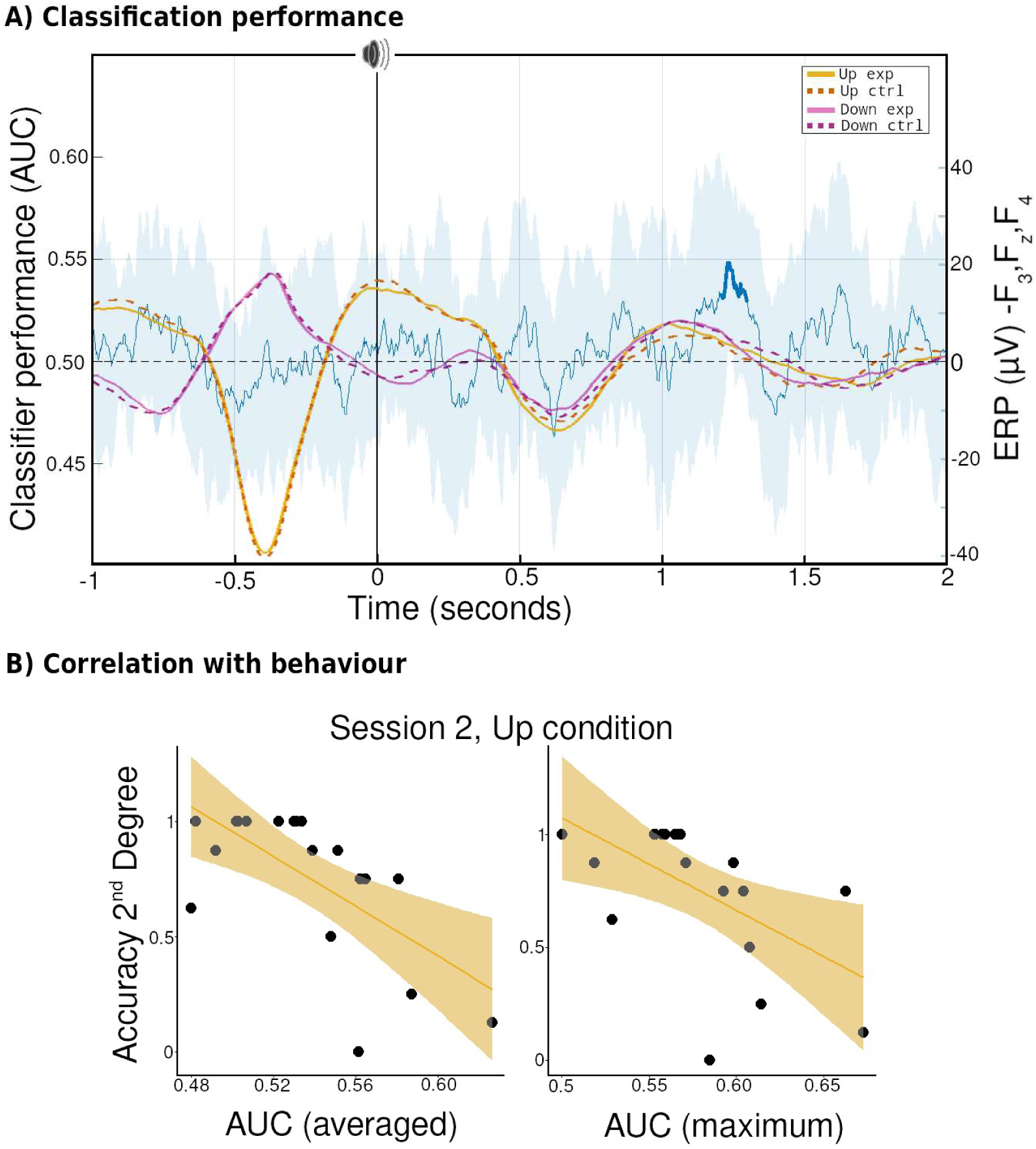
Classification. A) Up-Down classification performance. SVM classifier (blue line) presented a statistically significant cluster (over chance level) centered around 1.3s (thicker blue line). On top are superimposed the grand average ERPs value (mean of F3, Fz and F4 channels) for Up (yellowish colors) and Down (purple colors), with solid lines representing experimental sounds and dotted lines control sounds. B) Correlations between the SVM (AUC) mean -left-and peak performance -right-of the significant cluster for the Up condition with the behavioral accuracy of the second degree of difference-inference pair of the Up condition. Shadow areas represent 95% confident intervals.

To better understand the relationship between reactivation and consolidation, we performed a series of correlations between classification performance within the significant cluster (mean and peak) and the behavioral metrics described above. This revealed a negative correlation (R=-0.65, p_corrected_=0.022) between mean classification (SVM+AUC combination) and behavioral performance on 2^nd^ degree inference pairs in Session 2, figure 4B. This fits well with the fact that only 2^nd^ degree pairs in the Up condition presented a significant difference in overnight improvement when compared with the control condition for Session 2, and similarly only Up showed between condition differences in the degree of separation of inference pairs. The correlation was constant across all tested machine learning algorithms, see table SM5.1. No other correlation with any behavioral metric was significant (all p>0.05 before correction). However, the fact that this correlation is negative, e.g., the better the algorithm can classify the worse participants performed on that particular pair is perhaps surprising. Similar negative correlations have been reported, e.g. between classification performance and behavioral metrics (Abdellahi et al. 2021). One possible hypothesis is “good” performers try to fit the control sound into the previously learned hierarchy. That is, when they hear the Control sound, they may reactivate the cued hierarchy again, making it more difficult for the classifiers to differentiate between control from learned sounds.

## Discussion

Transitive inference is a key cognitive ability and a hallmark of deductive reasoning (Vasconcelos 2008). By showing that TMR in SWS strengthens such inferential thinking, our data support the idea that memory reactivation is important for more than the mere strengthening of memories which has been investigated by the bulk of the TMR literature (Hu et al. 2020). It is also involved in more complex integration and restructuring processes (Lewis and Durrant 2011). Furthermore, our findings show that such integration can be intentionally boosted through an external intervention, an observation which may be important for the development of cognitive enhancers targeting this kind of thinking. The fact that our manipulation had a stronger effect on the more distant 2^nd^ degree inferences is exciting in that it suggests the power of reactivation to assist in the integration of quite distant pieces of information.

### Phase of the slow oscillations and memory

The concept of an optimal phase or window for TMR stimulation stems from the fact that depolarising Up-states are more likely to activate larger groups of neurons synchronously (Vyazovskiy et al. 2009) and to drive thalamocortical spindles and sharp-wave ripples in the hippocampus (Batterink, Creery, and Paller 2016), thus facilitating hippocampal memory reactivation. This could explain why our classifiers only detect memory reactivation after Up stimulation, and why we found next-morning behavioural benefit only after stimulation in this phase. Our findings are in keeping with recent work showing that Up phase stimulation elicits classifiable reactivation and boosts memory performance, while Down phase stimulation does not (H.-V. V. Ngo and Staresina 2022) as well as work showing that Up phase TMR more effectively promotes emotional updating than stimulation at other SO phases (Xia et al. 2023). Thus, our results join a growing literature showing that Up phase TMR has a stronger impact on neural processing.

### TMR time course

The question of how long sleep related impacts on memory lasts is increasingly topical (Cordi and Rasch 2021; Pereira and Lewis 2020). One study reported that a TMR related reduction of implicit bias was retained after a week (Hu et al. 2020), while another study reported that TMR benefit to a serial reaction time task peaked after ten days (Rakowska et al. 2021). Here, accuracy for inference pairs was maintained two weeks after the manipulation for both Up and Control conditions (figure 2). However, performance in Down increased from below chance to above chance, reaching similar overall accuracy to the other two conditions after two-weeks (figure 2A-right). This improvement suggests that despite an initial inhibition due to TMR in the Down phase, the neural representation was able to recover over time. It is possible that subsequent nights of sleep without any manipulation may have allowed spontaneous reactivation of the memories, and that this occurred equally for all conditions. While Up and Control conditions, which had reactivated successfully in the first night, derived no benefit from this additional reactivation, the Down condition did benefit, and the associated consolidation allowed the Down-stimulated hierarchy to essentially catch up with the other hierarchies. Additionally, the next morning test in Session 2, which included presentation of inference pairs, may also have helped to trigger subsequent reactivation from which Down condition benefited.

## Summary

Our results show that the complex process of making indirect inferences can be facilitated by cued reactivation during slow wave sleep. Importantly, this was only true for the most distant inference pairs, and only Up phase cueing was effective. Importantly, these effects were not transitory, but instead lasted for the full two-week period examined here. These results provide strong support for the idea that memory reactivation in sleep is important for high-level qualitative changes in memory, such as integration and relational memory. Our findings also hold promise for the use of sleep-based interventions to drive improvement in such complex memory, and its application to real-word tasks.

## Acknowledgments

We would like to thank Elena Schmidt, Ralph Andrews and Duarte Pereira for contributing to the data collection, Dominic Carr for helping with the sleep scoring and the NAPs lab in general. This work was funded by the ERC Consolidator grant SolutionSleep, 681607 to PL.

## Supplementary materials

**Figure 1:**
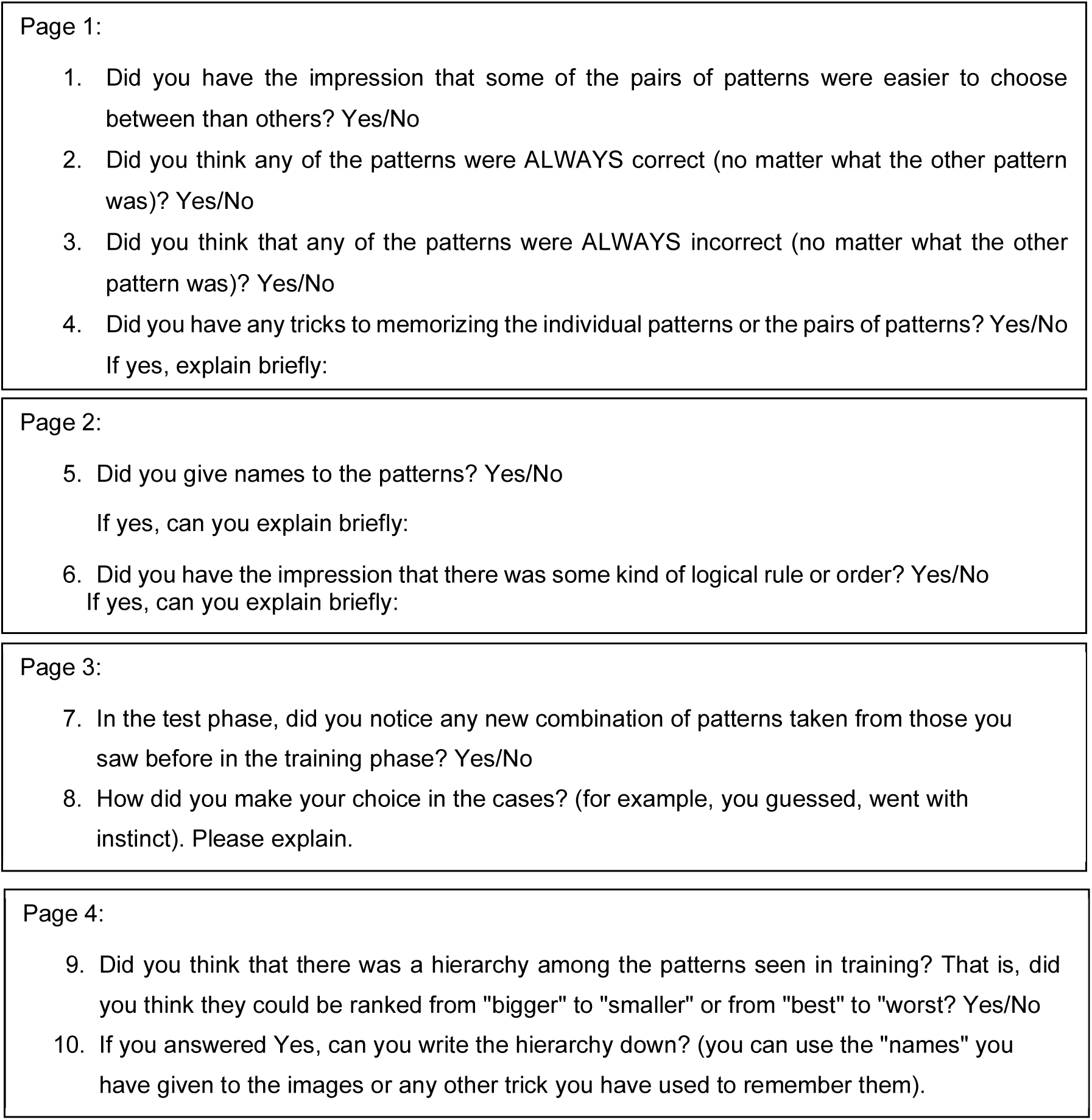
Questionnaire. Ten questions were asked at the end of the experiment dived into 4 pages as shown below:

**Figure 2:** Sound-Image association. Participants first learned the sound-image association to 90% criteria to increase chances that the overnight stimulation would trigger the associated image (see figure 1(A)). They were then tested on these associations but still received feedback to reinforce the learning. In the morning and during session 3 (2-weeks follow-up) they performed the same test but without feedback. A RM-ANOVA was performed to assess any difference in performance across sessions. Accuracy remained almost constantly at ceiling level for the three sessions (M:0.96 SE:0.004, M:0.97 SE:0.004, M:0.96 SE:0.004 respectively) with non-significant differences among them (smallest p=0.55).

**Figure 2-1:**
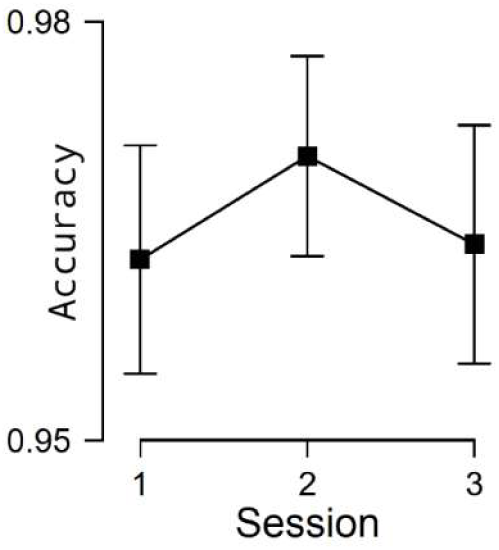
Sound-Image association performance. Accuracy for each session (evening, next morning and 2-weeks follow up). Horizontal bars represent 95 % confident intervals. No statistically significant differences were found.

**Figure 2-2:**
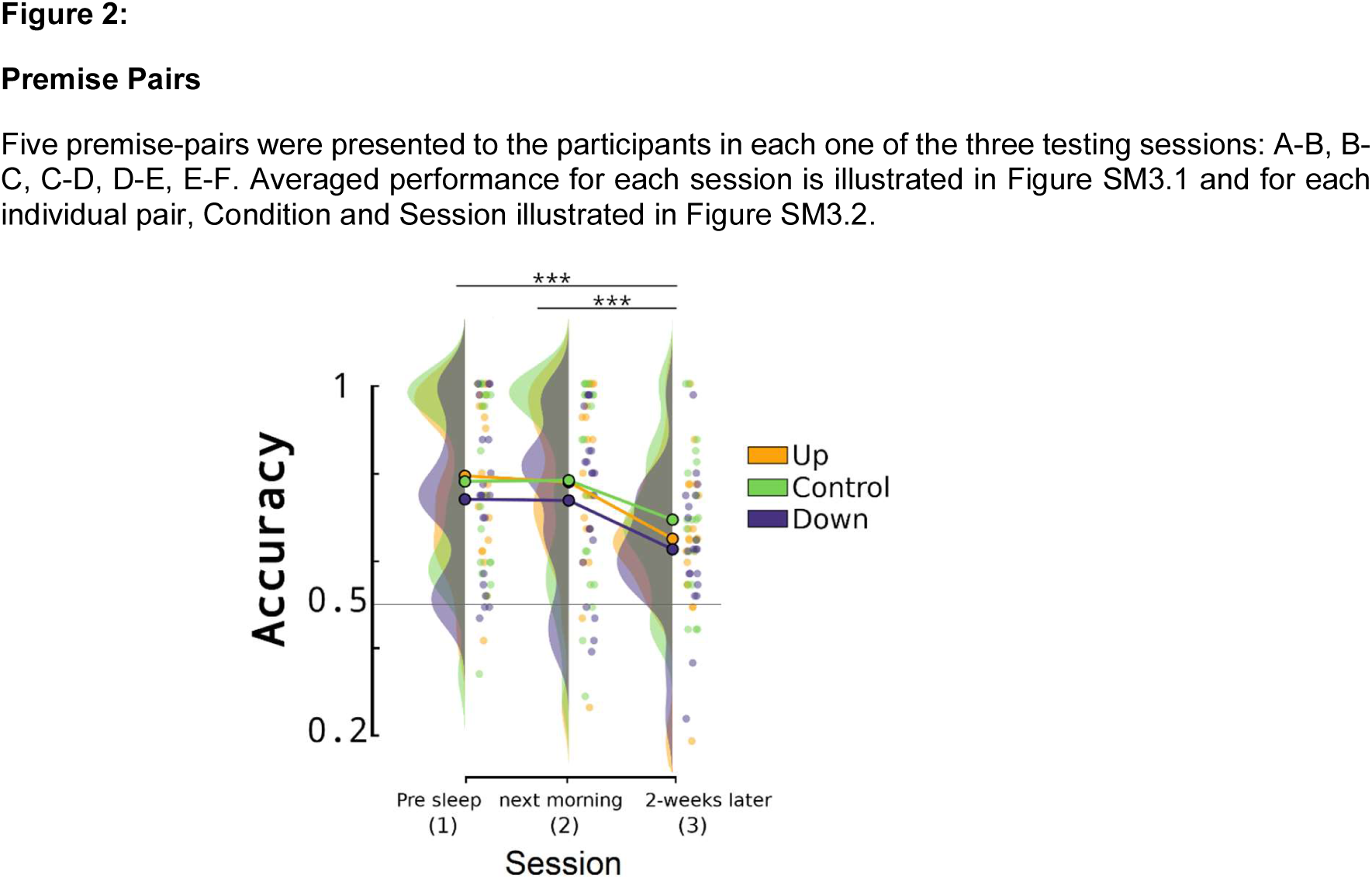
Premise pair accuracy results for each Session (x-axis) and for each one of the three conditions: Up (yellow), Control (green) and Down (purple). Statistically significant differences are indicated as: *p<0.05, ** p<0.01, ***p<0.001.

**Table 2-1:** Number of times each stimulus (F: Faces, O: objects, S: scenes) is used for each one of the three TMR conditions: Up, Down, Control.

| Stimuli | Up | Down | Control |
| --- | --- | --- | --- |
| F | 6 | 8 | 6 |
| O | 8 | 5 | 9 |
| S | 8 | 7 | 5 |

**Table 2-2:** Premise pair performance for each session (columns) and condition (rows). Accuracy is indicated as: number of trials, mean accuracy and standard error of the mean respectively. (*) Indicates statistically significant difference from chance level (50%).

|  | Session 1 |  |  | Session 2 |  |  | Session 3 |  |  |
| --- | --- | --- | --- | --- | --- | --- | --- | --- | --- |
|  | n | Mean | SE | n | Mean | SE | n | Mean | SE |
| Down | 390 | 0.747* | 0.018 | 390 | 0.747* | 0.019 | 340 | 0.629* | 0.022 |
| Control | 390 | 0.792* | 0.018 | 390 | 0.796* | 0.018 | 340 | 0.696* | 0.021 |
| Up | 390 | 0.796* | 0.018 | 390 | 0.782* | 0.018 | 340 | 0.653* | 0.022 |

**Figure 2-3:**
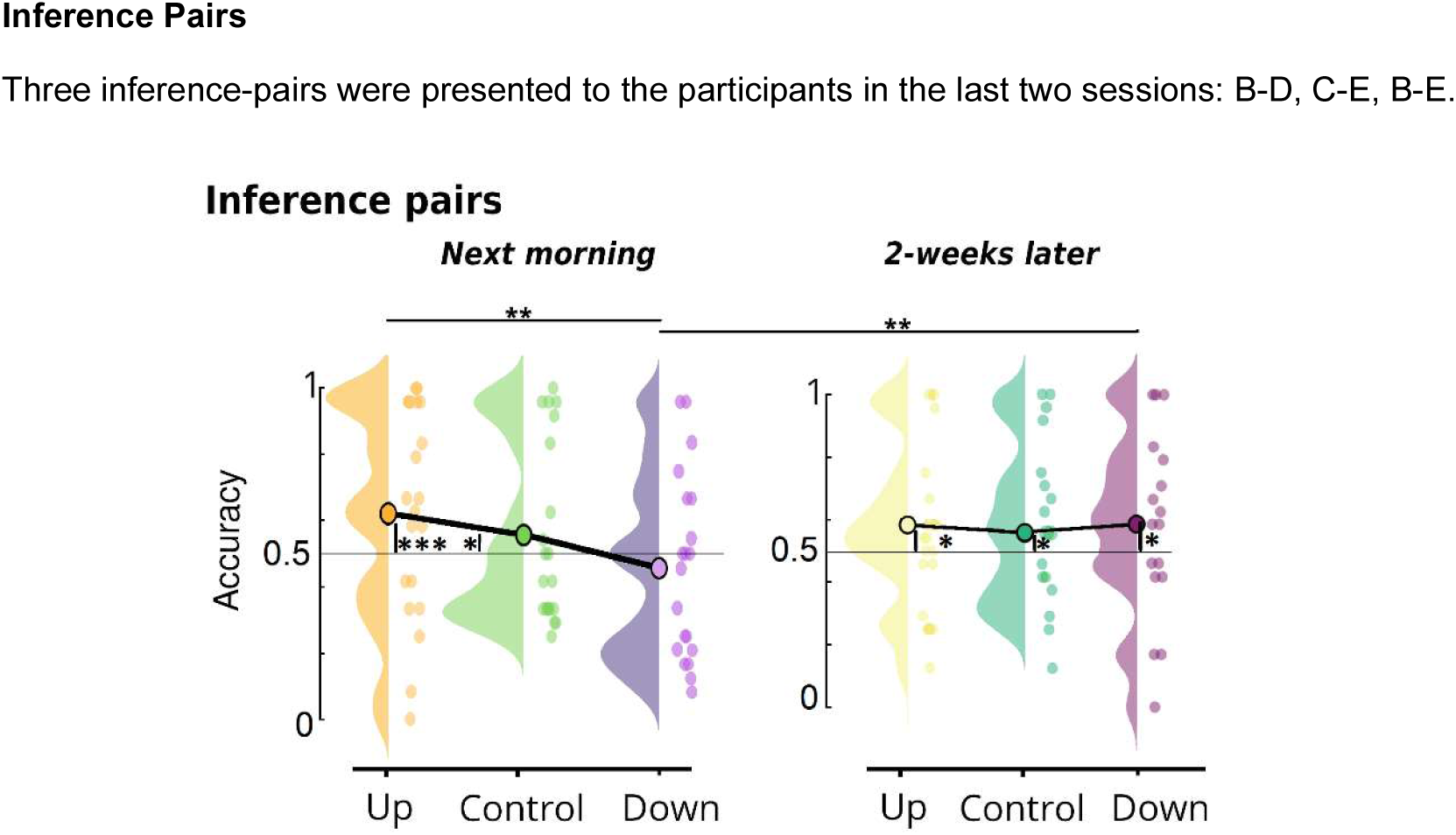
Inference pair performance for each condition: Up (yellow colours), Down (purple) and Control (greenish). Performance of averaged (1^st^ and 2^nd^ degree of separation) inference pairs for Session 2 (next morning) on the left and Session 3 (2-weeks later follow-up) on the right. Significant values from chance level are shown from the 50% line (chance level) for the first row (A).

**Table 2-3:** Inference pair performance for Overnight experiment. The first column represents the averaged pair performance, the other two the accuracy divided into the 1^st^ and 2^nd^ degree of separation. Accuracy is depicted for each session and condition of interest and indicated as: number of trials, mean accuracy and standard error of the mean respectively. (*) Indicates statistically significant difference from chance level (50%).

|  | Averaged |  |  | 1st Degree |  |  | 2nd Degree |  |  |
| --- | --- | --- | --- | --- | --- | --- | --- | --- | --- |
|  | n | Mean | SE | n | Mean | SE | n | Mean | SE |
| Session 2 |  |  |  |  |  |  |  |  |  |
| Down | 234 | 0.464 | 0.028 | 156 | 0.455 | 0.034 | 78 | 0.481 | 0.049 |
| Control | 234 | 0.560* | 0.027 | 156 | 0.574* | 0.032 | 78 | 0.532 | 0.049 |
| Up | 234 | 0.628* | 0.028 | 156 | 0.587* | 0.035 | 78 | 0.712* | 0.045 |
| Session 3 |  |  |  |  |  |  |  |  |  |
| Down | 204 | 0.586* | 0.029 | 136 | 0.577* | 0.036 | 68 | 0.603* | 0.050 |
| Control | 2048 | 0.559* | 0.029 | 136 | 0.555 | 0.036 | 68 | 0.566 | 0.050 |
| Up | 204 | 0.583* | 0.031 | 136 | 0.518 | 0.038 | 68 | 0.713* | 0.046 |

**Table 2-4:**
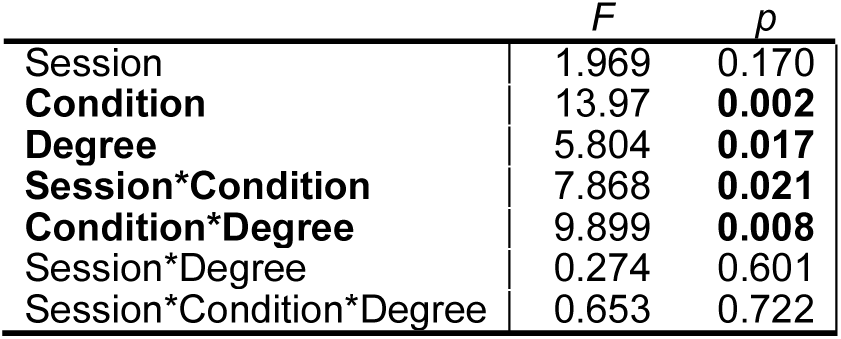
RM-ANOVA behavioral results with Session (2 and 3), Condition (Up, Down, Control) and Degree of separation (1^st^ and 2^nd^) as factors. Statistically significant results are highlighted in bold letters.

**Table 2-5:** Post-hoc Session*Condition interaction results. Statistically significant results are highlighted in bold letters.

| | $\Psi$ | *p* |
| --- | --- | --- |
| *Session 2* |  |  |
| Control vs Down | 2.25 | 0.214 |
| Control vs Up | -1.76 | 0.492 |
| **Down vs Up** | **-4.01** | **<0.001** |
| *Session 3* |  |  |
| Control vs Down | -0.707 | 0.981 |
| Control vs Up | -0.648 | 0.987 |
| Down vs Up | 0.059 | 0.997 |
| *Session 2 vs Session 3* |  |  |
| Control | -0.105 | 1.0 |
| **Down** | **-3.012** | **0.032** |
| Up | 0.924 | 0.94 |

**Table 2-6:**
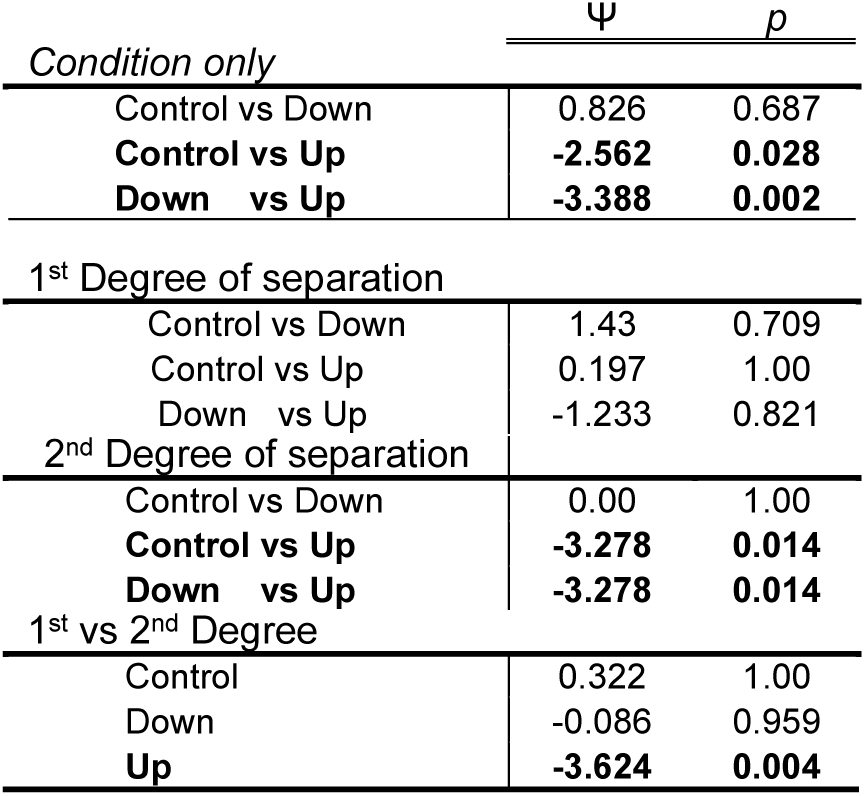
Post-hoc Degree*Condition interaction results. Statistically significant results are highlighted in bold letters.

**Table 2-7:** Statistical analysis per session for the 2^nd^ Degree of difference per Condition. Statistically significant results are highlighted in bold letters.

| 2<sup>nd</sup> Degree | Session 2 |  | Session 3 |  |
| --- | --- | --- | --- | --- |
| | $\Psi$ | $p$ | $\Psi$ | $p$ |
| Control vs Down | 0.077 | 0.880 | -0.531 | 0.856 |
| **Control vs Up** | **-0.416** | **0.027** | -2.124 | 0.087 |
| **Down vs Up** | **-0.494** | **0.006** | -1.593 | 0.251 |

**Figure 4.** **Classifiers.** We used two classification algorithms, SVM and LDA, with two different performance metrics, AUC (area under the curve) and ACC (accuracy) to distinguish between Experimental and Control sounds. Cluster statistics resulted in a consistent positive cluster for the Up (see table 4-1) condition but not significant clusters for the Down condition (see table 4-2). Results of the SVM classifier with AUC as performance metrics for both conditions are shown in Figure 4-1. Additionally, we performed correlations between the classification performance and the behavioral results for the Up condition taking both the mean and the peak of the significant classifier cluster. Results for the mean performance within the cluster can be seen in table 4-3 and results for the peak performance within the cluster in table 4-4.

**Table 4-1:**
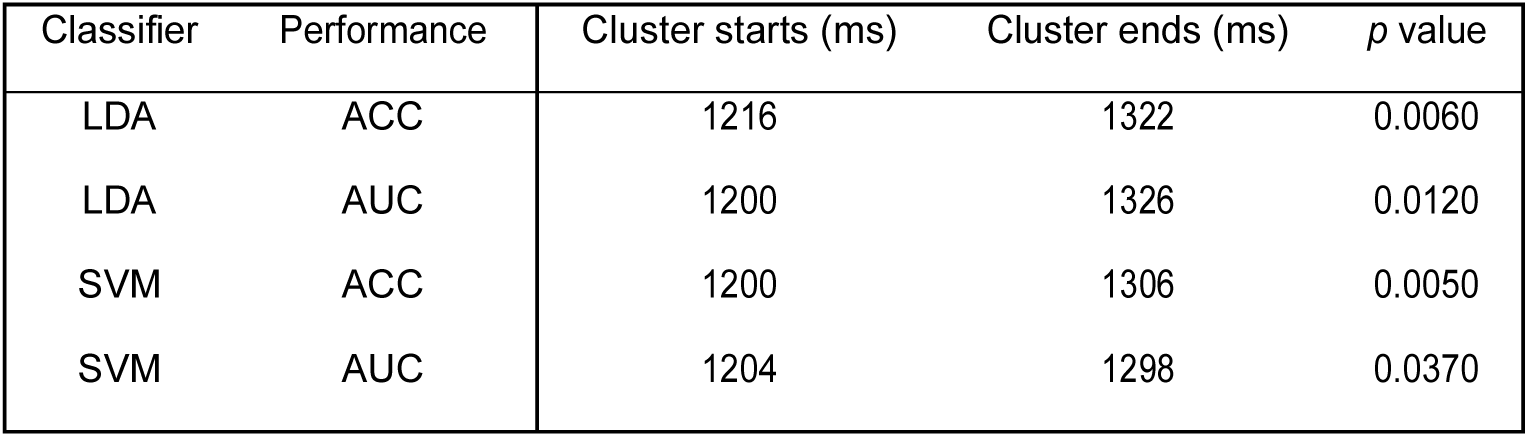
Significant cluster across different algorithms for Up condition

**Table 4-2:** Cluster statistic results across different algorithms for Down condition.

| Classifier | Performance | lowest p value |
| --- | --- | --- |
| LDA | ACC | 0.095 |
| LDA | AUC | 0.143 |
| SVM | ACC | 0.106 |
| SVM | AUC | 0.074 |

**Table 4-3:** Correlations results between classifier performance (mean within the significant cluster) and behavioral accuracy of the second degree-inference pairs for the Up condition.

| Classifier | Rho | $p$ original | $p$ corrected |
| --- | --- | --- | --- |
| LDA ACC | -0.59 | 0.006 | 0.028 |
| LDA AUC | -0.59 | 0.006 | 0.031 |
| SVM ACC | -0.64 | 0.002 | 0.027 |
| SVM AUC | -0.65 | 0.002 | 0.022 |

**Figure 4-1:**
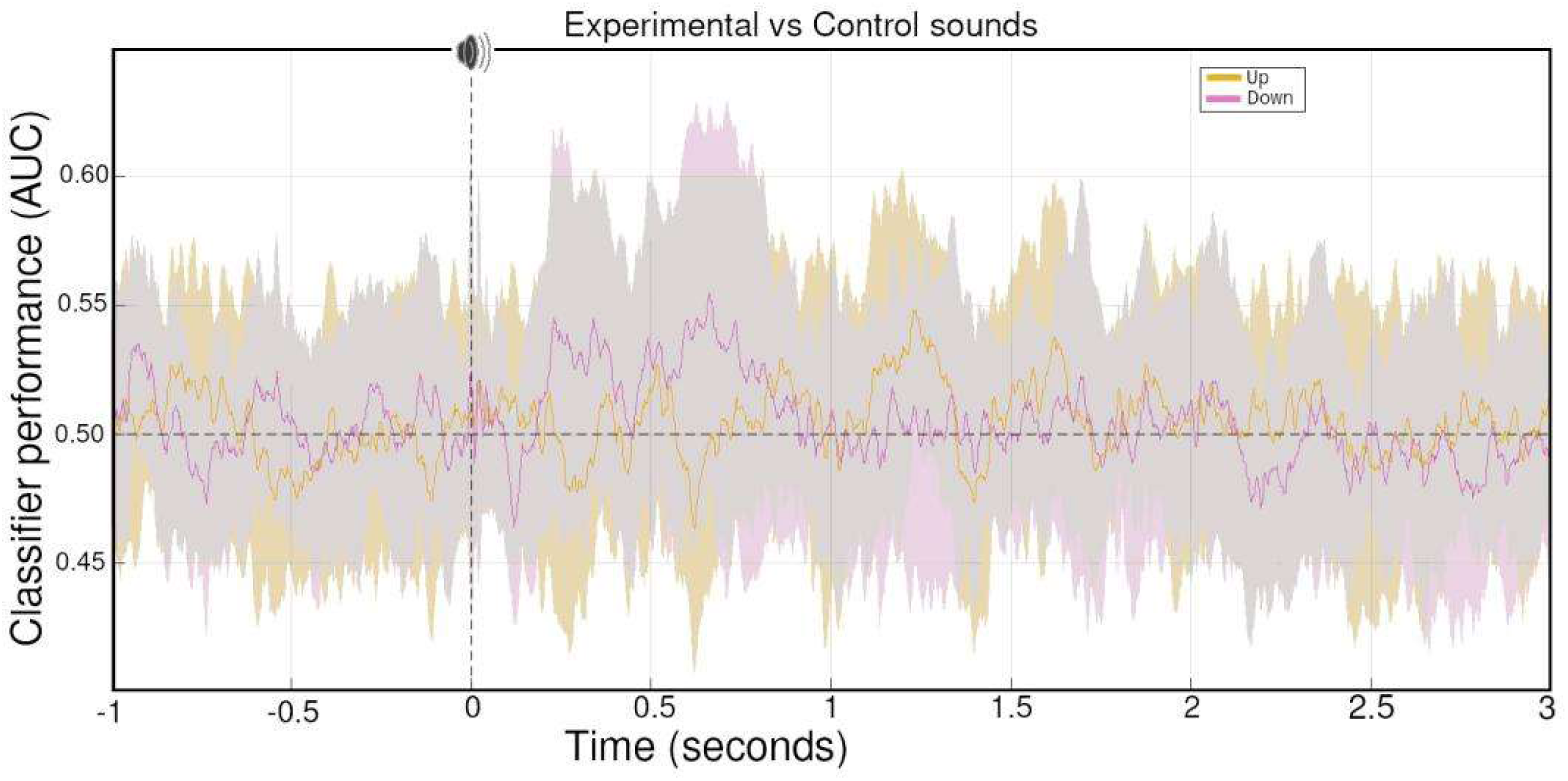
Grand average classifier results for Up (yellowish) and Down (purple) conditions using a SVM with AUC as performance metric. Shadow areas corresponding to the stand deviation across participants.

**Table 4-4:** Correlations results between classifier performance (peak of the significant cluster) and behavioral accuracy of the second degree-inference pairs for the Up condition.

| Classifier | Rho | $p$ original | $p$ corrected |
| --- | --- | --- | --- |
| LDA AUC | -0.49 | 0.039 | 0.063 |
| LDA ACC | -0.48 | 0.039 | 0.065 |
| SVM ACC | -0.57 | 0.012 | 0.042 |
| SVM AUC | -0.56 | 0.012 | 0.047 |

## Notes

### Competing Interest Statement

The authors have declared no competing interest.

### Summary of Updates

Updated analysis Corrected grammar errors

